# A robust and adaptive framework for interaction testing in quantitative traits between multiple genetic loci and exposure variables

**DOI:** 10.1101/2021.12.01.469907

**Authors:** Julian Hecker, Dmitry Prokopenko, Matthew Moll, Sanghun Lee, Wonji Kim, Dandi Qiao, Kirsten Voorhies, Woori Kim, Stijn Vansteelandt, Brian D. Hobbs, Michael H. Cho, Edwin K. Silverman, Sharon M. Lutz, Dawn L. DeMeo, Scott T. Weiss, Christoph Lange

**Affiliations:** Channing Division of Network Medicine, Brigham and Women’s Hospital, Boston, MA, USA; Genetics and Aging Unit and McCance Center for Brain Health, Department of Neurology, Massachusetts General Hospital, Boston, MA, USA; Division of Pulmonary and Critical Care Medicine, Brigham and Women’s Hospital, Boston, MA, USA; Department of Medical Consilience, Division of Medicine, Graduate School, Dankook University, South Korea; PRecisiOn Medicine Translational Research (PROMoTeR) Center, Department of Population Medicine, Harvard Pilgrim Health Care, Boston, MA, USA; Systems Biology and Computer Science Program, Ann Romney Center for Neurological Diseases, Department of Neurology, Brigham and Women’s Hospital, Boston MA, USA; Department of Applied Mathematics, Computer Science and Statistics, Ghent University, Gent, Belgium; Department of Medical Statistics, London School of Hygiene and Tropical Medicine, London, UK; Department of Biostatistics, Harvard T.H. Chan School of Public Health, Boston, MA, USA; Harvard Medical School, Boston, MA, USA

## Abstract

The identification and understanding of gene-environment interactions can provide insights into the pathways and mechanisms underlying complex diseases. However, testing for gene-environment interaction remains a challenge since statistical power is often limited, the specification of environmental effects is nontrivial, and such misspecifications can lead to false positive findings. To address the lack of statistical power, recent methods aim to identify interactions on an aggregated level using, for example, polygenic risk scores. While this strategy increases power to detect interactions, identifying contributing key genes and pathways is difficult based on these global results.

Here, we propose RITSS (Robust Interaction Testing using Sample Splitting), a gene-environment interaction testing framework for quantitative traits that is based on sample splitting and robust test statistics. RITSS can incorporate multiple genetic variants and/or multiple environmental factors. Using sample splitting, a screening step enables the selection and combination of potential interactions into scores with improved interpretability, based on the user’s unrestricted choices for statistical/machine learning approaches. In the testing step, the application of robust test statistics minimizes the susceptibility of the results to main effect misspecifications.

Using extensive simulation studies, we demonstrate that RITSS controls the type 1 error rate in a wide range of scenarios. In an application to lung function phenotypes and human height in the UK Biobank, RITSS identified genome-wide significant interactions with subcomponents of genetic risk scores. While the contributing single variant interactions are moderate, our analysis results indicate interesting interaction patterns that result in strong aggregated signals that provide further insights into gene-environment interaction mechanisms.

## Introduction

Genome-wide association studies (GWAS) have identified thousands of genetic variants that are associated with complex diseases/phenotypes (MacArthur *et al*., 2017). However, the effect of a genetic variant on a complex phenotype or disease can be modified by environmental exposures (Hunter, 2005; Khoury, 2017). The knowledge about an interaction between environmental exposure (e.g., tobacco smoke or occupational exposure) and genetic variants can provide insights into the underlying pathways and disease mechanisms (Thomas, 2010).

Most of the methodological approaches for gene-environment interaction testing have focused on scenarios in which a single genetic variant and a single environmental variable are tested for potential interaction. Methods have been developed for case-control data, case-only data, and quantitative trait studies, summarized in detail by Gauderman et al. (Gauderman *et al*., 2017).

Since most gene-environment interaction effects are expected to be small and statistical power is consequently limited (Murcray *et al*., 2011), more efficient approaches were proposed. This includes so-called screening statistics that aim to prioritize genetic variants to reduce the multiple testing problem (Dai *et al*., 2012; Gauderman *et al*., 2010, 2013; Hsu *et al*., 2012; Kooperberg and Leblanc, 2008; Murcray *et al*., 2011, 2009; Paré *et al*., 2010; Zhang *et al*., 2016). Also, researchers proposed to aggregate genetic information in a genomic region in set-based tests to increase power (Jiao *et al*., 2013; Liu *et al*., 2016; Lin *et al*., 2013; Tzeng *et al*., 2011; Zhao *et al*., 2015; Lin *et al*., 2016; Su *et al*., 2017; Jiao *et al*., 2015; Kim *et al*., 2019). Recent approaches to increase power utilize mixed models or reaction models and incorporate multiple environmental variables to derive a combination of environmental factors that modifies the genetic effect (Moore *et al*., 2019; Ni *et al*., 2019; Dahl *et al*., 2020; Kerin and Marchini, 2020; Wang *et al*., 2020).

Also, current data analyses observed that genetic risk scores, weighted sums of genetic risk variants, interact with specific environmental factors in psychiatric (Peyrot *et al*., 2018) and cardiovascular diseases (Hindy *et al*., 2018). In addition, Aschard et al. observed an interaction between a genetic risk score for FEV_1_/FVC and ever-smoking status (Aschard *et al*., 2017). Related, Kim et al. also observed highly significant interactions between genome-wide polygenic risk scores for lung function and smoking variables (Kim *et al*., 2021). While detecting significant interactions on this aggregated level is encouraging, identification of individual genes and pathways is challenging based on such results.

Besides the power limitation, another caveat of gene-environment interaction testing is that the misspecification of the marginal main effect models can lead to inflated type 1 error rates and false positive findings in interaction testing (Sun *et al*., 2018; Zhang *et al*., 2020). This is especially problematic for non-binary and/or continuous environmental factors where the implicit linearity assumption that is commonly used in the model might not be correct. One example is pack-years of smoking, where it is not straightforward to assume that every additional pack-year has the same, constant effect on the outcome.

In this communication, we propose RITSS, a robust and flexible framework for gene-environment interaction testing with quantitative traits. The general idea is to derive an interaction score comprised of the (weighted) sum of individual genetic variant / environmental factor pairs, such that the combination of these signals increases the power to detect an interaction while maintaining the biological interpretability of the results. To provide accurate statistical results, our approach utilizes a sample splitting strategy and test statistics that are robust against misspecifications of the main effects. The general form of RITSS allows the incorporation of user-specified screening/learning approaches in the construction of candidate interaction scores while providing valid statistical inference without restrictive assumptions. In extensive simulation studies, we demonstrate the robustness of RITSS in various realistic scenarios. We applied RITSS to lung function phenotypes and human height in the UK Biobank, incorporating sex and smoking exposure information. Our analyses identified highly consistent interaction patterns across sets of genetic variants that result in statistically significant interactions with subcomponents of genetic risk scores, while the interaction effects at the single variant level are moderate.

## Methods

We use the following notation: for study subject *i*, we denote the quantitative trait of interest by *Y*_*i*_, the *m*-dimensional genotype information by *X*_*i*_, and the *d*-dimensional environmental information by *E*_*i*_. Also, we define an additional *p*-dimensional covariate vector by *Z*_*i*_ that includes, for example, age, study indicators, and genetic principal components (Price *et al*., 2006). We assume the following model

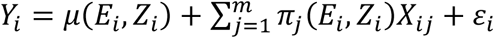

where *E*[*ε*_*i*_|*X*_*i*_, *Z*_*i*_, *E*_*i*_] = 0. The unknown function *μ* describes the main effect of the environmental factors *E*_*i*_ and other covariates *Z*_*i*_. The genetic contribution of variant *j* is modeled by *π*_*j*_(*E*_*i*_, *Z*_*i*_)*X*_*ij*_, where *π*_*j*_ is an unknown function. The null hypothesis of no gene-environment interaction is described by 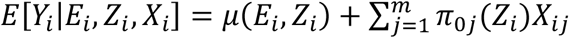, where *π*_0*j*_ is an unknown function only depending on *Z*_*i*_. Therefore, we implicitly assume the absence of gene-gene-interactions but allow for interactions between the genetic variants and the covariates *Z*_*i*_ under the null hypothesis. Testing for gene-environment interaction in quantitative traits between a single variant *j* and a single environmental factor *l* is often underpowered. On the other side, testing for interaction between an environmental factor *l* and a genetic risk score that combines genetic information across loci can have more power if the directions of the interaction effects are in line with the summation in the genetic risk score. However, the interpretation of interactions with such dense scores can be difficult. Our approach, RITSS, therefore aims to derive interaction scores of the form *U*_*i*_ = ∑_*j*_ ∑_*l*_ *π*_*jl*_*X*_*ij*_*E*_*il*_ that combine signals while maintaining biological interpretability of the overall interaction result by keeping the number of involved factors of moderate size. As combining potential interaction signals into a score in a data-adaptive way and testing the score needs, in general, to be performed in independent parts of the data, RITSS utilizes a sample splitting approach. Another challenge in gene-environment interaction testing is the test statistic itself. The standard interaction score test, for example, for the pair consisting of variant *j* and environmental factor *l*, is based on the following quantity:

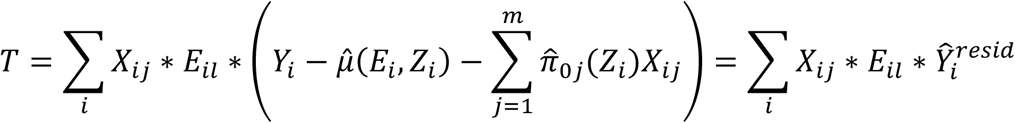

Under the null hypothesis of no gene-environment interaction between variant *j* and environmental factor *l*, the test statistic *T* needs to have expectation 0 to provide a valid test. This is the case if approximately 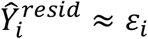, meaning that 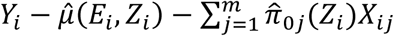 captures the main effects reasonably well. However, if, for example, the environmental main effect of environmental factor *l* is slightly mis specified, 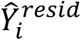 can have some residual dependency on *E*_*il*_. Then, in general, *T* does not have expectation 0 under the null hypothesis since the residual term can be correlated with *X*_*ij*_ * *E*_*il*_. The observation that mis specified main effects lead to inflated interaction type 1 error rates has been described in the recent literature (Sun *et al*., 2018; Zhang *et al*., 2020). Furthermore, when the number of genetic variants, environmental factors, and covariates (i.e., *m, d*, and *p*) increases, residual dependency is more likely to occur since convergence rates of main effect models will become slower due to the increased complexity.

Therefore, we utilize an approach based on test statistics that improve robustness against such misspecifications (Vansteelandt *et al*., 2008). Denoting the variable *U*_*i*_ as the interaction score, the general idea is to replace 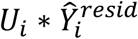 by 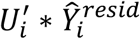 where 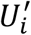 is a projection of *U*_*i*_ that is orthogonal to the environmental and genetic main effects.

Thereby, in the presence of slight misspecifications or residual dependencies of the main effects, the test statistic based on 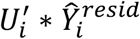 has expectation 0 under the null hypothesis of no gene-environment interaction. The form of the test statistic will also allow alternating the roles of training and testing subsamples to use the full sample size for testing. The details are described in the next section.

### RITSS

We now describe the algorithm behind RITSS. The algorithm is also visualized in Figure 1.

**Figure 1.**
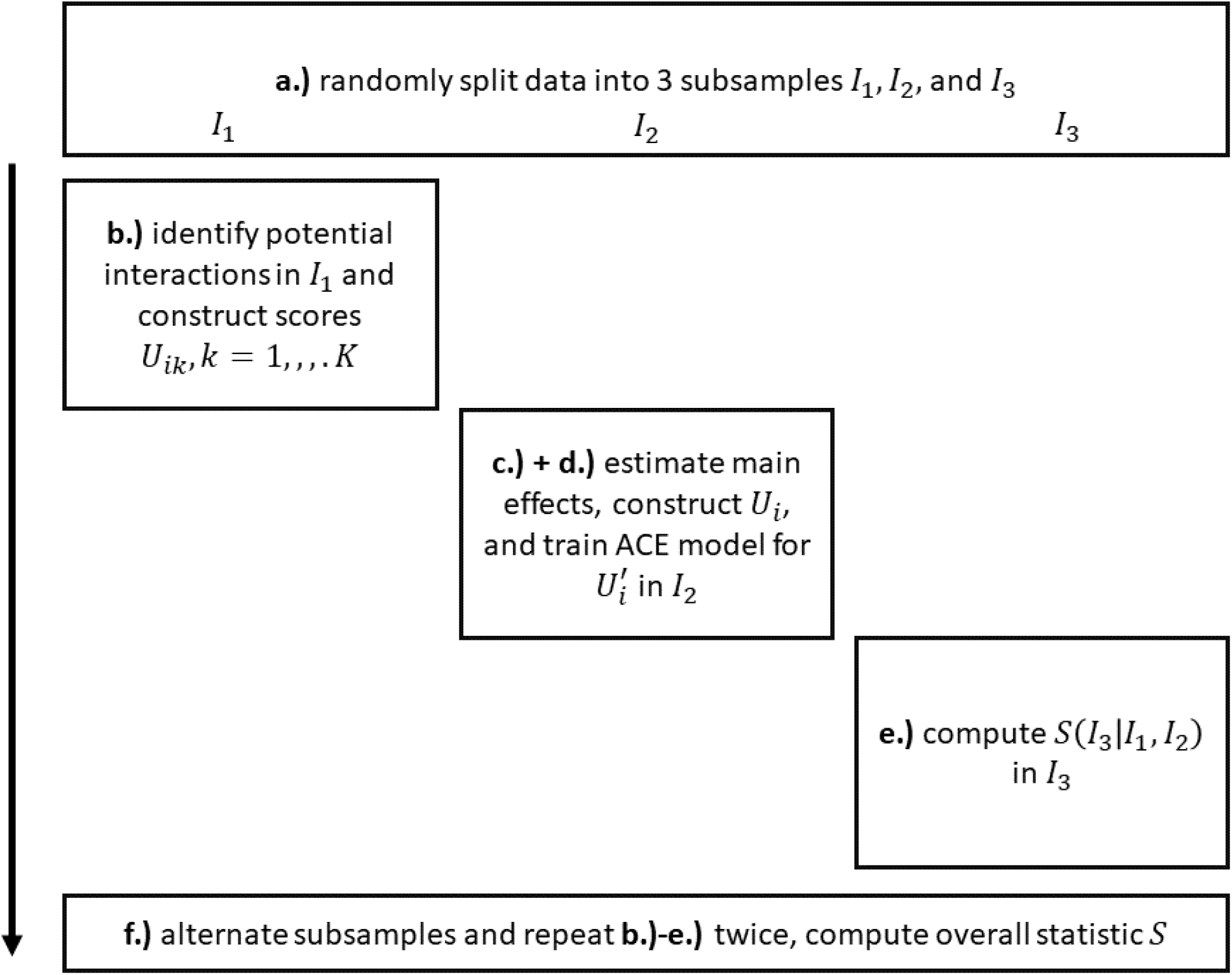
Visualization of the algorithm underlying our RITSS framework.

### Algorithm

#### a.) Sample splitting

First, we split the data (*Y*_*i*_, *X*_*i*_, *E*_*i*_, *Z*_*i*_) randomly into three approximately equally sized subsamples *I*_1_, *I*_2_, and *I*_3_.

#### b.) Discovery step using *I*_1_

Using *I*_1_ data, we construct potential interaction scores of the form *U*_*ik*_ = ∑_*j*_ ∑_*l*_ *π*_*jlk*_*X*_*ij*_*E*_*il*_, *k* = 1, …, *K* that aim to combine signals. Here, the number of scores, *K* can be pre-specified or adaptively chosen. The information about which pairs of genetic/environmental factors are included and the determination of the corresponding weights *π*_*jlk*_ is based on the user’s choice of statistical/machine learning-based approaches to model the phenotypic data. Examples include LASSO, Bayesian shrinkage, and best subset regressions (Tibshirani, 1996; Castillo *et al*., 2015; Bertsimas *et al*., 2016). Usage of *K* > 1 and therefore partitioning into different scores could reflect certainty about the corresponding interaction effects. An example is described in the application. It is important to note that, as the screening step generates scores that are tested in different parts of the data, the screening method can be based on arbitrary methods, and no statistical rigor is required, but suitable specifications will increase the overall power of our approach.

#### c.) Estimation of main effects and further filtering using *I*_2_

Using the first half of the *I*_2_ data, we fit the interaction scores *U*_*ik*_, *k* = 1, …, *K*, derived in Step b.), as predictors in a linear model while incorporating flexible models for the main effects 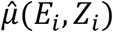 and 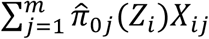. These flexible main effect models are modeled using linear terms with higher-order terms in combination with penalization (LASSO-based techniques (Tibshirani, 1996)). The motivation here is that we want to estimate the main effects as accurately as possible while fitting the proposed interaction scores to check their significance. Since *I*_2_ is an independent part of the data, the gene-environment interaction scores *U*_*ik*_ can be considered as fixed covariates here and significance of these scores in *I*_2_ provides further evidence of interaction. We select all or a subset of scores (for example, based on the significance of their regression coefficients) and combine them (potentially with estimated weights) into a single interaction score *U*_*i*_. Then, the estimated main effects 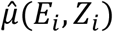 and 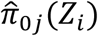 are extracted and are used to compute 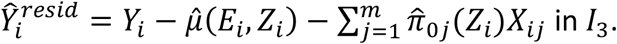.

#### d.) Deriving 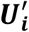in *I*_3_

As outlined above, standard interaction testing would consider a test statistic based on 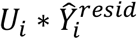. To obtain a robust test statistic, we replace this with 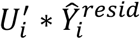, where 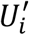 is a projection of *U*_*i*_ that is orthogonal to the environmental and genetic main effects. This projection can be obtained by the alternating conditional expectation (ACE) algorithm (Breiman and Friedman, 1985; Vansteelandt *et al*., 2008). If gene and environment are independent, closed-form solutions are available. We apply this algorithm to the second half of the *I*_2_ data. Finally, based on the trained model, we compute *U*_*i*_′ in *I*_3_. The details of the implementation of ACE are described in Appendix B.

#### e.) Computing test statistic in *I*_3_

Based on these objects, we compute our interaction test statistic in *I*_3_:

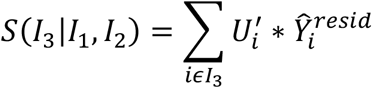

#### f.) Alternating subsamples

We alternate the roles of the subsamples and repeat steps b.)-e.) twice, then compute the overall statistic

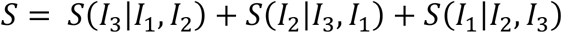

### Properties of *S*

The test statistic *S* in combination with the described procedure a.)-f.) to obtain the underlying objects has several important advantages. Under mild regularity conditions, the statistic *S* is asymptotically normal, the sub statistics are asymptotically independent, and the variance of *S* can be estimated based on a simple empirical variance estimator. By estimating a flexible model for the main effects in the computation of 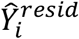 and the application of the ACE to obtain 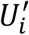 the interaction test based on *S* is robust against (residual) misspecifications of the main effect models. Due to the sample splitting approach, we do not have to assume strict regularity conditions for the estimation of machine learning models and still can use the full sample size for testing (Vansteelandt *et al*., 2008; Dukes *et al*., 2021).

It is important to note that the interaction scores that are tested in the three sub statistics are not identical in general, since they are constructed from different parts of the data. The idea is that the overall interaction RITSS p-value provides evidence for the presence of interaction based on these adaptive scores. Additional insights can be obtained by examining the overlap between the three scores.

The RITSS approach is flexible and allows for different targeted applications and the incorporation of arbitrary statistical learning approaches in the screening step. We will describe a specific application of RITSS in the next section and investigate its performance in simulations and in the application to the UK Biobank.

### Simulation studies and UK Biobank analysis

In this section, we discuss a specific implementation of RITSS that aims to identify interactions between a single environment factor of interest and subcomponents of genetic risk scores. In simulation studies, we demonstrate the validity and robustness of this approach in different scenarios. We also apply the approach to the UK Biobank to analyze interactions in lung function and height.

### Specific implementation of RITSS

Steps a.), d.), e.), and f.) are performed as described in the Methods section.

#### b.) Discovery step using *I*_1_

Using *I*_1_ data, we first fit the model 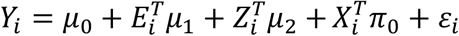. Then, we construct the matrix 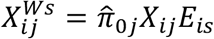 where *s* is the index of the environmental factor of interest. The matrix 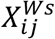 describes genotype-environment products weighted by the corresponding estimated genetic main effect. Next, we regress out *X*_*i*_, *E*_*i*_, and *Z*_*i*_ from *Y*_*i*_ and 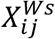 (for each variant *j*) and denote the resulting residuals by 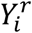 and 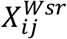. The variant-wise variances of 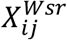, denoted by *υ*_*j*_, differ typically substantially due to the different weights of genotypic effects and minor allele frequencies. We keep only variants whose variance is larger than 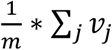 and denote the resulting number of variants by *S*_*max*._ Based on these objects, we split *I*_1_ randomly in two equally sized parts and perform approximate best subset selection (Bertsimas *et al*., 2016) in each part with different subset sizes, where the corresponding other part is used as testing data. Best subset here corresponds to the best subset of variants, and we do not utilize the effect sizes per variant inferred in the best subset regression. Instead, we use the summation over the corresponding variants in 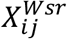, describing the genetic risk score multiplied by the environmental factor. The size of the subsets is increased between 5 and *S*_*max*_, in steps of 5. We select the best subsets in both parts of the data in terms of association with 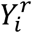 in the test data and check the overlap between these subsets to create two different scores. Denote by *b*_1_ the set of overlapping variants (contained in both best subsets), and by *b*_2_ the set of non-overlapping variants that were only in one best subset. The first score is then defined by 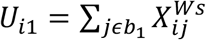, and the second by 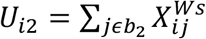 (and they can be computed in *I*_2_ and *I*_3_ using exactly these weights). Overall, these two scores represent genetic risk scores multiplied by the environmental factor of interest, identified in combination with the main effect working model 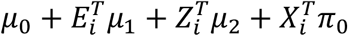.

#### c.) Estimation of main effects and further filtering using *I*_2_

Using the first half of *I*_2_ data, we fit the model 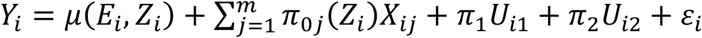, where *μ*(*E*_*i*_, *Z*_*i*_) and 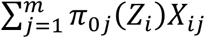 are realized using higher-order and product terms in combination with penalization (LASSO) and cross-validation. Based on the fitted model, we obtain 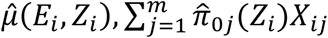 and the overall interaction score *U*_*i*_ = *U*_*i*1_ + *c*_2_*U*_*i*2_, where *c*_2_ is set to 1, if the corresponding regression coefficient p-value is below 0.05, otherwise *c*_2_ = 0. All genetic variants that are included in *U*_*i*_, and therefore tested with the sub statistic *S*(*I*_3_|*I*_1_, *I*_2_), are denoted by the set *m*(*I*_3_|*I*_1_, *I*_2_).

### Simulations

We performed extensive simulations to demonstrate the validity and robustness of RITSS based on the described implementation. The goal of the simulation studies is to show that the approach provides controlled type 1 error rates in a variety of scenarios, under the null hypothesis of no gene-environment interactions.

### Simulation settings

Our simulations are based on different specifications of the model

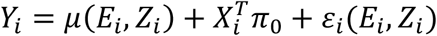

with *E*[*ε*_*i*_(*E*_*i*_, *Z*_*i*_)|*X*_*i*_, *Z*_*i*_, *E*_*i*_] = 0. The scenarios are based on a combination of the following elements:

1. Population stratification (PS): *Z*_*i*_ influences *E*_*i*_, *X*_*i*_, and *Y*_*i*_
2. Gene-environment correlation (GEC): *X*_*i*_ influences *E*_*i*_
3. Mis specified environmental main effects (MEM): 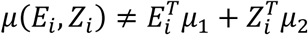 for all vectors *μ*_1_, *μ*_2_.
4. Non-normal errors (NNE): Error distribution of *ε*_*i*_ not normal
5. Heteroscedasticity (HE): Variance of *ε*_*i*_ depends on *E*_*i*_.

Point 3.) impacts the construction of interaction scores in the screening step b.) where a simple linear working model for main effects is assumed. Overall, 32 scenarios can be generated as combinations of the presence or absence of these five factors. For the simulations with non-normal errors, we used the mean centered and standardized FEV_1_/FVC data from the UK Biobank as the random error to investigate robustness against this error distribution. In addition, we investigated the question if the type 1 error rate is inflated in the case where the SNPs for the analysis were selected based on GWAS results in the same dataset, i.e., based on their GWAS association p-values (notated as SELECT yes/no). Incorporating this factor, we, therefore, simulated a total of 64 scenarios. In all simulations, we set a sample size of *n* = 30,000, *m* = 100 SNPs, *d* = 5 environmental factors *E*_*i*_, and *p* = 5 principal components *Z*_*i*_. All simulation results were evaluated based on 1,000 replicates. The specific details of how the presence or absence of the six factors PS yes/no, GEC yes/no, MEM yes/no, NNE yes/no, HE yes/no, and SELECT yes/no were implemented are described in Appendix A.

### Comparison with linear regression

To demonstrate the importance of the robust test statistic in our approach, we also included a standard regression analysis as a comparison. Here, we use the interaction score *U*_*i*_, derived by the steps a.) to d.), but instead of using the robust statistic *S*(*I*_3_|*I*_1_, *I*_2_) and 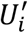 to test the interaction in *I*_3_, we incorporate *U*_*i*_ as a covariate in the regression model

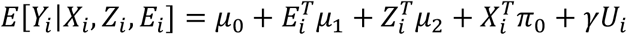

in *I*_3_ and test for interaction based on the coefficient *γ*. We denote this approach by REG. We also incorporated REG-robust, where the corresponding standard errors are computed based on a robust sandwich-variance-estimator. Since we only consider type 1 error rate studies and not power results, the fact that REG/REG-robust use *I*_3_ data only, whereas RITSS used the whole dataset, is not crucial.

## Results

We report the distribution of p-values for the three methods RITSS, REG, and REG-robust in all 64 scenarios in Figures 2-5 and Supplementary Figures 1-4. Figures 2-5 visualize the quantile-quantile-plots (qq-plots) for SELECT:no, partitioned according to the four combinations of GEC yes/no and MEM yes/no. Accordingly, Supplementary Figures 1-4 report the corresponding qq-plots for SELECT:yes.

**Figure 2:**
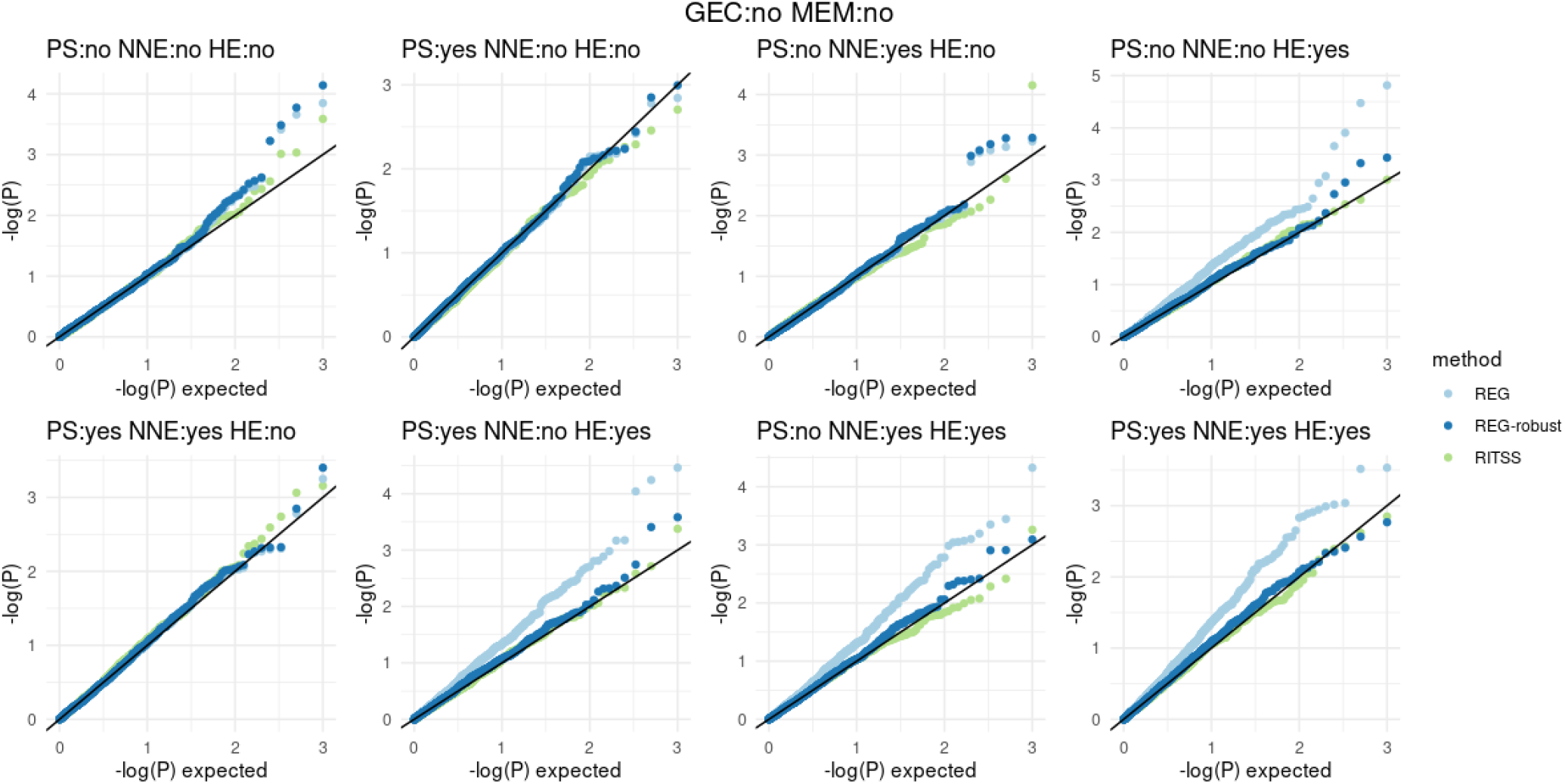
Quantile-quantile-plots for RITSS, REG, and REG-robust in the 8 scenarios with SELECT:no, GEC:no, and MEM:no. All results based on 1,000 replicates. GEC: gene-environment correlation, MEM: mis specified environmental main effect, PS: population stratification, NNE: non-normal errors, HE: heteroscedasticity.

Figure 2 shows the qq-plots for GEC:no and MEM:no. All methods control the type 1 error rates, except REG in the sub scenarios with HE:yes. This is expected because the standard error estimation in REG assumes homoscedastic errors. The results in Figure 3 (GEC:yes and MEM:no) are similar. In the case of GEC:no and MEM:yes (Figure 4), we see that REG is inflated in all sub scenarios since the mis specified environmental main effect introduces heteroscedastic errors (Sun *et al*., 2018). REG-robust is able to provide controlled type 1 error rates due to the robust variance estimation. RITSS also controls the type 1 error rates in these scenarios. In Figure 5 (GEC:yes and MEM:yes), REG and REG-robust are highly inflated in all 8 sub scenarios, while RITSS still remains valid. The explanation is that the presence of GEC and MEM leads to constructions of *U*_*i*_ that seemingly capture interactions but are just the implication of the misspecifications of the main effects in the screening step and the regression test model. Therefore, whereas the robustness property of RITSS still ensures valid results (due to the more flexible main effect estimation in step c.) and the projected score in step d.)), the regression-based approaches become inflated. The results for the SELECT:yes scenarios (Supplementary Figure 1-4) are similar to the corresponding results for SELECT:no. Overall, the simulations demonstrate the robustness of RITSS in a variety of different scenarios.

**Figure 3:**
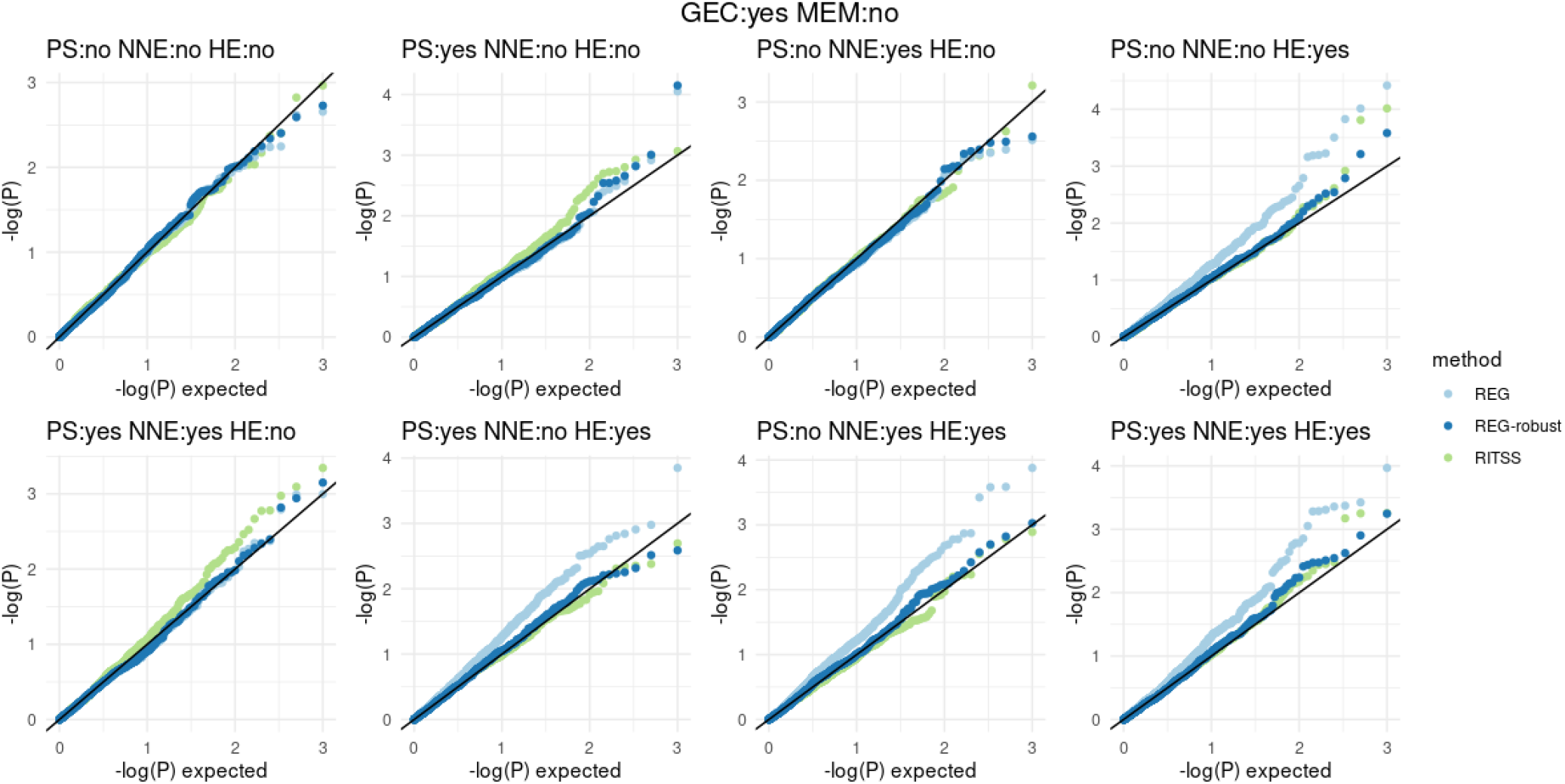
Quantile-quantile-plots for RITSS, REG, and REG-robust in the 8 scenarios with SELECT:no, GEC:yes, and MEM:no. All results based on 1,000 replicates. GEC: gene-environment correlation, MEM: mis specified environmental main effect, PS: population stratification, NNE: non-normal errors, HE: heteroscedasticity.

**Figure 4:**
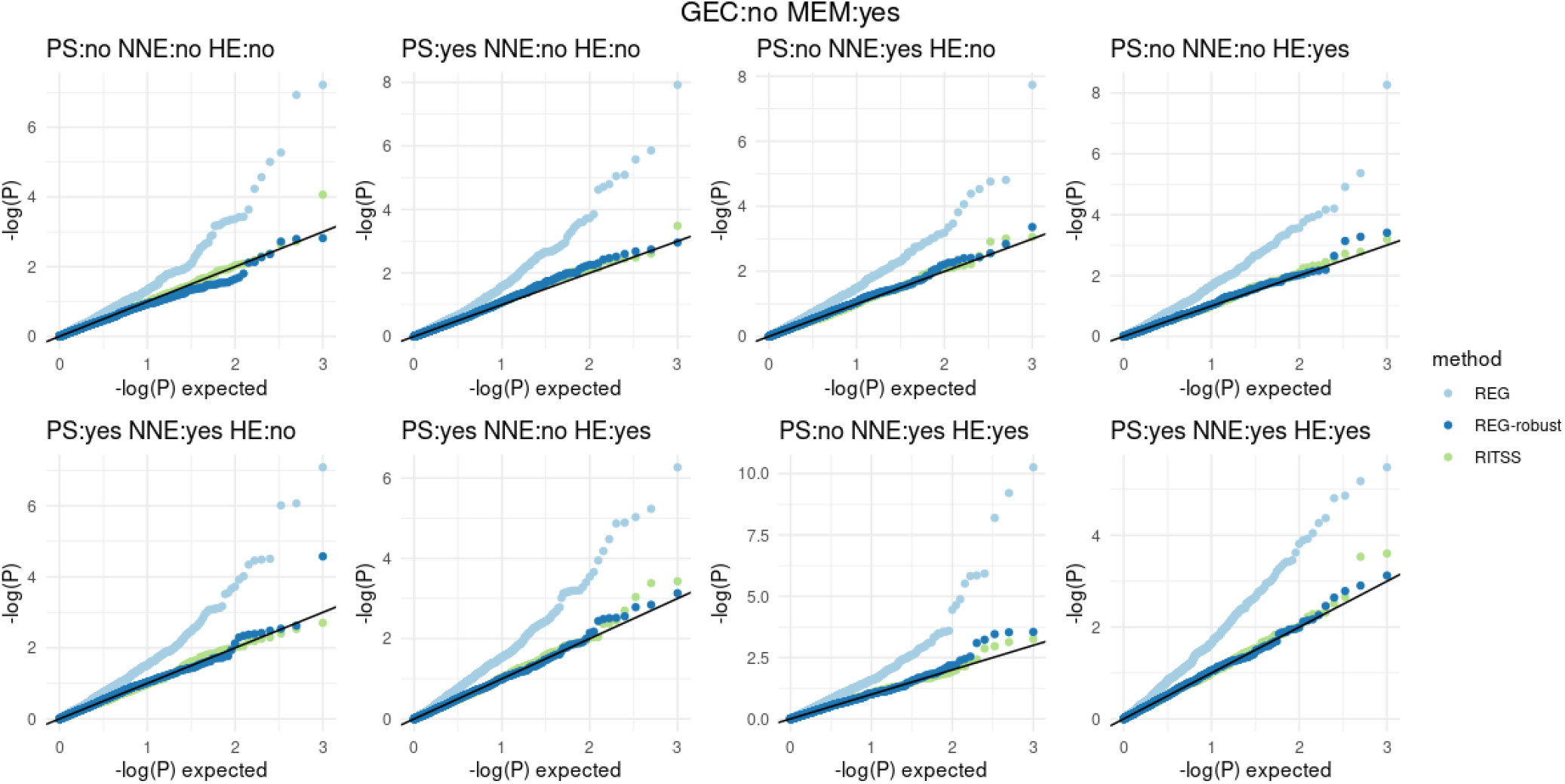
Quantile-quantile-plots for RITSS, REG, and REG-robust in the 8 scenarios with SELECT:no, GEC:no, and MEM:yes. All results based on 1,000 replicates. GEC: gene-environment correlation, MEM: mis specified environmental main effect, PS: population stratification, NNE: non-normal errors, HE: heteroscedasticity.

**Figure 5:**
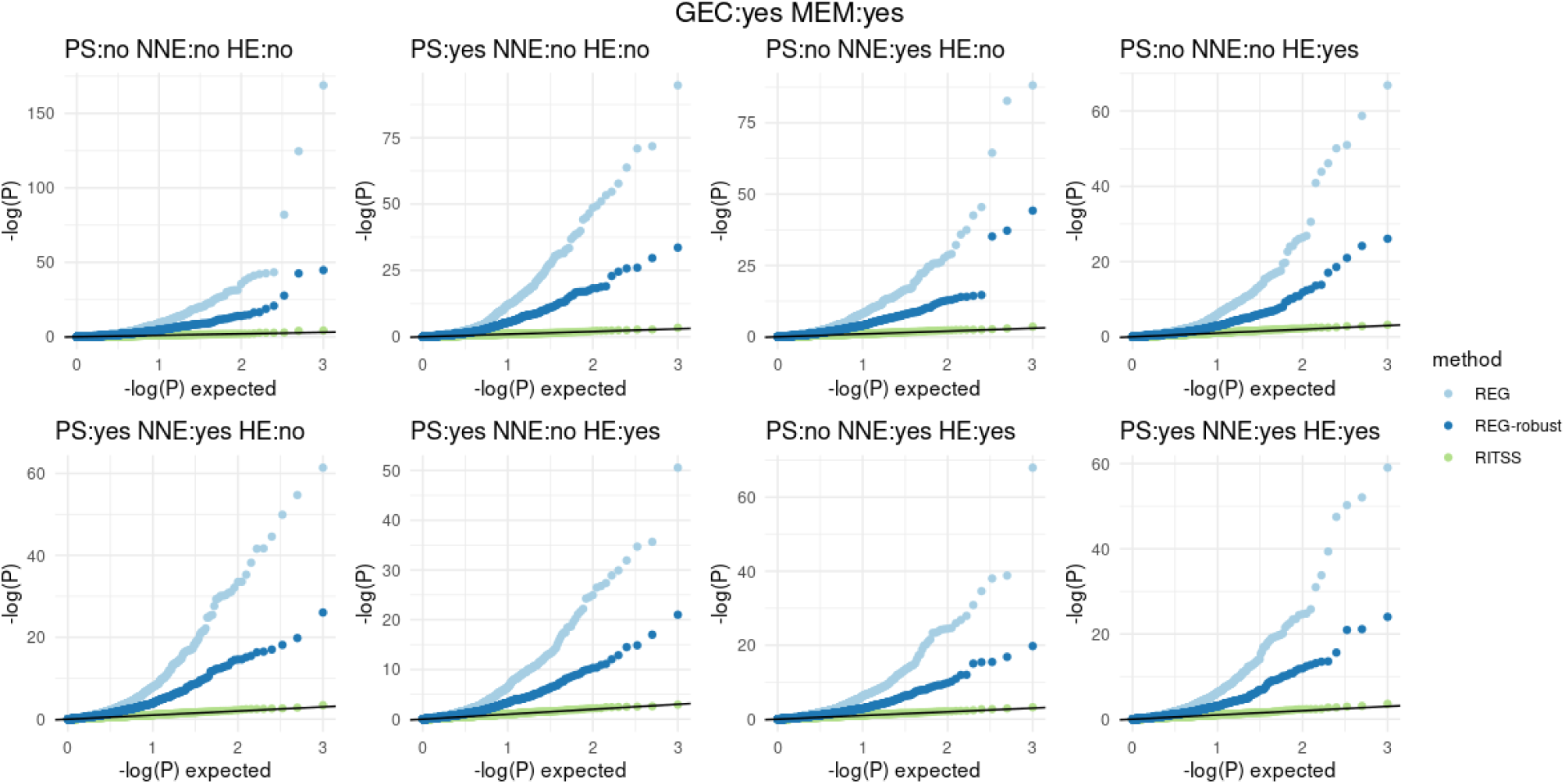
Quantile-quantile-plots for RITSS, REG, and REG-robust in the 8 scenarios with SELECT:no, GEC:yes, and MEM:yes. All results based on 1,000 replicates. GEC: gene-environment correlation, MEM: mis specified environmental main effect, PS: population stratification, NNE: non-normal errors, HE: heteroscedasticity.

In addition, in Figure 6, we plotted the qq-plot based on 3*64 pairwise correlation p-values between the z-scores of the sub statistics *S*(*I*_3_|*I*_1_, *I*_2_), *S*(*I*_2_|*I*_3_, *I*_1_), and *S*(*I*_1_|*I*_2_, *I*_3_) across all simulated scenarios. The results show that the sub statistics are indeed asymptotically independent, which is also reflected in the controlled type 1 error rates of the p-values that are based on the variance estimator that exploits the independence.

**Figure 6:**
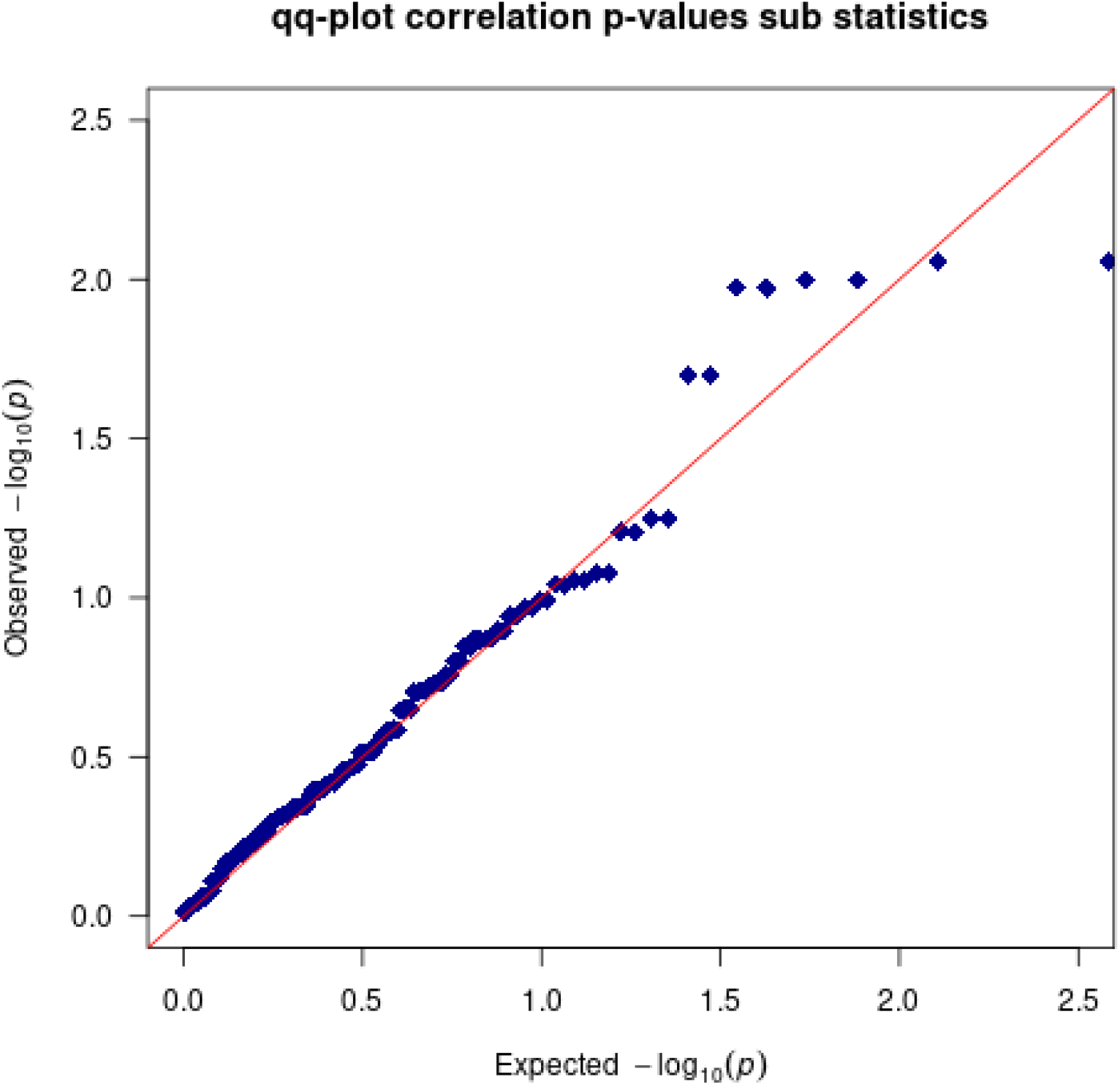
Quantile-quantile-plots of the 3*64 pairwise correlation p-values between the z-scores of the corresponding three sub statistics in the 64 simulated scenarios. The results are based on 1,000 replicates.

### UK Biobank analysis

We applied RITSS to the UK Biobank data to analyze gene-environment interactions for lung function (measured by forced expiratory volume in 1 second (FEV_1_), forced vital capacity (FVC) and the ratio FEV_1_/FVC) and height. Details about the study population as well as the extraction of genetic, environmental, and phenotypic data are described in Appendix C.

### Analysis setup

Table 1 contains the configurations of *Y*_*i*_, *X*_*i*_, *E*_*i*_, and *Z*_*i*_ for the four different analyses. Age and pack-years of smoking (P-Y-S) were mean centered before computing the squared variable. Sex was coded as male=1 and female=0. For the lung function traits, we also included height as an interacting variable. We note that, given this specific sex coding, height and sex are (strongly) positively correlated. The lung function measurements were analyzed on the original scale and not transformed, as robustness of the approach against non-normal errors was demonstrated in the simulations. After quality control, we split the 254,053 samples (European ancestry) in two parts: 180,000 randomly selected samples for the main analysis using RITSS and 74,053 samples that are not analyzed by RITSS and serve as an independent set to validate the analysis results of RITSS. For the analysis, we applied the RITSS as described above. Each of the factors in *E*_*i*_ was tested for interaction, resulting in a total of 16 tests. In Supplementary Figure 5, we plotted the estimated densities of the standardized residuals after adjusting for *X*_*i*_, *E*_*i*_, and *Z*_*i*_ for the selected traits. The density plots are based on the 180,000 samples in the main analysis.

**Table 1.** Analysis configurations and number of genetic variants incorporated. P-Y-S: pack-years of smoking, E-S: ever-smoking, PCs: genetic principal components. FEV_1_: forced expiratory volume in 1 second, FVC: forced vital capacity (FVC). Pack-years of smoking and age were centered before computing the squared variable.

| $Y_i$ | $X_i$ | $E_i$ | $Z_i$ |
| --- | --- | --- | --- |
| FEV<sub>1</sub> | 383 SNPs | sex, height, P-Y-S, E-S, P-Y-S<sup>2</sup> | age, age<sup>2</sup>, PCs 1-10, genotyping-array |
| FVC | 638 SNPs | sex, height, P-Y-S, E-S, P-Y-S<sup>2</sup> | age, age<sup>2</sup>, PCs 1-10, genotyping-array |
| FEV<sub>1</sub>/FVC | 885 SNPs | sex, height, P-Y-S, E-S, P-Y-S<sup>2</sup> | age, age<sup>2</sup>, PCs 1-10, genotyping-array |
| height | 3368 SNPs | sex | age, age<sup>2</sup>, PCs 1-10, genotyping-array |

### Results

Table 2 contains the results of our UK Biobank RITSS analysis using 180,000 samples. For each of the four traits, we observed a significant interaction between sex and subcomponents of the genetic risk score. For the lung function traits, we also observed similar interaction findings with height. We investigated this in more detail in the validation analysis below.

**Table 2.** Results of the interaction testing using RITSS in the UK Biobank. The tested environmental factor is denoted by *E*_*is*_. |*m*| is the number of total SNPs in the analysis, |*m*(*I*_3_|*I*_1_, *I*_2_)|, |*m*(*I*_2_|*I*_3_, *I*_1_)| and |*m*(*I*_1_|*I*_2_, *I*_3_)| denote the number of SNPs in the three sub statistics. |*m*_3_| and |*m*_2_| are the number of SNPs that are shared by all three and exactly two interaction scores, respectively.

| $Y_i$ | $E_{is}$ | RITSS p-value | $\vert m\vert $ | $\vert m(I_3\vert I_1, I_2)\vert $ | $\vert m(I_2\vert I_3, I_1)\vert $ | $\vert m(I_1\vert I_2, I_3)\vert $ | $\vert m_3\vert $ | $\vert m_2\vert $ |
| --- | --- | --- | --- | --- | --- | --- | --- | --- |
| FEV<sub>1</sub> | sex | 5.834e-34 | 383 | 145 | 130 | 137 | 72 | 67 |
| FEV<sub>1</sub> | height | 9.121e-14 | 383 | 145 | 131 | 132 | 68 | 70 |
| FEV<sub>1</sub> | P-Y-S | 3.012e-01 | 383 | 7 | 12 | 6 | 0 | 1 |
| FEV<sub>1</sub> | E-S | 4.220e-01 | 383 | 8 | 5 | 25 | 0 | 4 |
| FEV<sub>1</sub> | P-Y-S<sup>2</sup> | 3.853e-01 | 383 | 10 | 23 | 9 | 0 | 6 |
| FVC | sex | 2.670e-22 | 638 | 248 | 215 | 243 | 61 | 137 |
| FVC | height | 5.241e-18 | 638 | 248 | 249 | 221 | 56 | 152 |
| FVC | P-Y-S | 9.923e-03 | 638 | 9 | 6 | 149 | 0 | 5 |
| FVC | E-S | 2.945e-01 | 638 | 61 | 10 | 202 | 1 | 24 |
| FVC | P-Y-S<sup>2</sup> | 6.344e-01 | 638 | 39 | 15 | 61 | 0 | 6 |
| FEV<sub>1</sub>/FVC | sex | 3.392e-24 | 885 | 300 | 303 | 288 | 110 | 156 |
| FEV<sub>1</sub>/FVC | height | 4.112e-17 | 885 | 294 | 171 | 262 | 70 | 119 |
| FEV<sub>1</sub>/FVC | P-Y-S | 4.687e-05 | 885 | 291 | 295 | 232 | 89 | 143 |
| FEV<sub>1</sub>/FVC | E-S | 1.232e-05 | 885 | 287 | 23 | 288 | 10 | 151 |
| FEV<sub>1</sub>/FVC | P-Y-S<sup>2</sup> | 6.809e-01 | 885 | 297 | 175 | 98 | 32 | 107 |
| height | sex | 2.619e-14 | 3368 | 1013 | 895 | 618 | 275 | 427 |

Furthermore, in line with previous results in the literature, we observed a significant interaction with ever-smoking and pack-years of smoking for FEV_1_/FVC (Kim *et al*., 2021; Aschard *et al*., 2017).

As outlined above, the three interaction scores tested in each of the three sub-statistics can consist of different genetic variants, due to the independent screening in different parts of the data. These sets of genetic variants are denoted by *m*(*I*_3_|*I*_1_, *I*_2_), *m*(*I*_2_|*I*_3_, *I*_1_), and *m*(*I*_1_|*I*_2_, *I*_3_). Furthermore, we denote the set of genetic variants that are shared by all three or exactly two sub statistics by *m*_3_ and *m*_2_, respectively. Table 2 reports the number of genetic variants in these sets.

Based on the genetic variants in *m*_2_ and *m*_3_, we performed an additional validation analysis using the remaining and independent 74,053 samples. We considered all interactions in Table 2 that were significant after Bonferroni correction adjusting for 16 tests at an overall significance level of *α* = 0.05. For this validation analysis, we used a standard regression interaction test while fitting *X*_*i*_, *E*_*i*_, and *Z*_*i*_. We tested two interaction scores, the first based on the variants in the corresponding *m*_3_ set and the second based on *m*_2_. The interaction p-values were evaluated based on the model-based standard errors as well as standard errors obtained from robust sandwich estimators. Here, the genotype main effect estimates for the genetic risk scores were estimated based on the 180,000 samples in the main analysis.

The results of our validation analysis are described in Table 3. The results show that almost all findings replicate except the *m*_3_ based E-S interaction for FEV_1_/FVC. We also note that the effect directions were consistent between the main analysis and validation analysis.

**Table 3.** Results of the validation analysis in the 74,053 remaining samples in the UK Biobank. The two sets *m*_3_ and *m*_2_ were identified in the corresponding main analysis (Table 2). The interaction p-values are based on a standard regression analysis, the robust p-values are based on standard errors obtained by a robust sandwich variance estimator.

| $Y_i$ | $E_{is}$ | p-value $m_3$ | robust p-value $m_3$ | $\vert m_3\vert $ | p-value $m_2$ | robust p-value $m_2$ | $\vert m_2\vert $ |
| --- | --- | --- | --- | --- | --- | --- | --- |
| FEV<sub>1</sub> | sex | 8.161e-22 | 5.072e-11 | 72 | 9.625e-05 | 1.961e-04 | 67 |
| FEV<sub>1</sub> | height | 5.162e-10 | 4.431e-09 | 68 | 2.593e-06 | 9.029e-06 | 70 |
| FVC | sex | 1.732e-05 | 2.962e-05 | 61 | 1.149e-09 | 6.074e-09 | 137 |
| FVC | height | 3.938e-06 | 1.142e-05 | 56 | 3.686e-09 | 6.702e-08 | 152 |
| FEV<sub>1</sub>/FVC | sex | 9.325e-11 | 2.510e-10 | 110 | 2.660e-04 | 3.178e-04 | 156 |
| FEV<sub>1</sub>/FVC | height | 4.754e-03 | 4.912e-03 | 70 | 7.130e-06 | 1.076e-05 | 119 |
| FEV<sub>1</sub>/FVC | P-Y-S | 4.995e-07 | 8.929e-05 | 89 | 2.197e-02 | 8.743e-02 | 143 |
| FEV<sub>1</sub>/FVC | E-S | 7.570e-01 | 7.648e-01 | 10 | 1.374e-03 | 2.403e-03 | 151 |
| height | sex | 8.720e-05 | 1.051e-04 | 275 | 8.327e-07 | 1.108e-06 | 427 |

To examine these findings in more detail, we examined the single variant interaction p-values for the genetic variants in the combined set *m*_3_/*m*_2_ with the respective environmental factor in the validation dataset. This analysis was also based on a linear regression that fits *X*_*i*_, *E*_*i*_, and *Z*_*i*_, and the respective interaction term. Table 4 lists the number of genome-wide significant single variant interactions, i.e., *p* < 5 * 10^−8^, as well as the number of significant single variant interactions using a Bonferroni correction adjusting for the number of variants in the combined set *m*_3_/*m*_2_ at a significance level of *α* = 0.05. Both numbers are either 0 or very small, for all trait/environmental factor combinations. This suggests that the interaction effects at the single variant level are small. However, we make an interesting observation by investigating the empirical distribution of effect directions across the single variant interaction tests. Table 4 provides these empirical distributions of effect direction for a.) the unweighted interaction terms *X*_*ij*_ * *E*_*il*_ and b.) the weighted terms 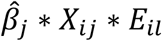, where 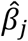 is the estimated genetic effect based on the main analysis dataset. In addition, we compared the interaction p-value distributions according to the effect direction of the weighted interaction terms. As we can see by the enrichment of positive effects, the weighting by the genetic main effect, as it is applied in the computation of the genetic risk score, “aligns” the interaction effect directions across the genetic variants. Furthermore, the p-values for weighted interaction terms with positive effect show higher inflation. This indicates that RITSS identified genetic variants whose interaction signals are weak at the single variant level but when summed up in the genetic risk score align to a significant interaction. Since we used a sex coding of male=1 and female=0, this suggests that the genetic effects in males and females across these variants have the same direction, but the magnitudes of effects in males are slightly larger. Related, we note that the genetic variants in *m*_3_/*m*_2_ between the sex and height interactions with lung functions highly overlap, which is in line with the observation that height and sex are strongly positively correlated. However, if we test the height interaction separately in males and females in the validation dataset, the signal is strongly diminished. Therefore, we hypothesize that the observed interactions are a result of sex-differential effects, but further analyses are required to disentangle the exact mechanisms. In this context, we also note that, related to recent results regarding so-called participation biases and their impact on sex-related analyses (Pirastu *et al*., 2021), the genetic variants in *m*_3_/*m*_2_ across the three lung function phenotypes and height were not associated with sex. Detailed information about the selected variants in the gene-by-sex interactions for lung function and height and the corresponding closest genes (obtained by the GWAS catalogue data) can be found in Supplementary Tables 1-4.

**Table 4.** Results of the validation analysis in the 74,053 remaining samples in the UK Biobank. The two sets *m*_3_ and *m*_2_ were identified in the corresponding main analysis (Table 2). All variants in *m*_3_/*m*_2_ were tested for interaction with the respective environmental factor *E*_*is*_. In this context, gws. interaction refers to a single variant interaction p-value *p* < 5 * 10^−8^. Similar, bfs. Interaction corresponds to *p* < 0.05/(|*m*_3_|+|*m*_2_|). The last columns report the empirical distribution of effect directions in these tests and the corresponding 5% quantile of p-values. In the weighted analysis, the genotype/env.factor product term was multiplied by the estimated genetic main effect (obtained from independent data).

| $Y_i$ | $E_{is}$ | $\vert m_3\vert +\vert m_2\vert $ | No. gws<br>interactions | No. bfs.<br>interactions | Distribution effect<br>directions unweighted<br>-/+ | Distribution<br>effect directions<br>weighted -/+ | 5% quantile<br>interaction p-values in<br>weighted -/+ |
| --- | --- | --- | --- | --- | --- | --- | --- |
| FEV<sub>1</sub> | sex | 139 | 0 | 1 | 0.57/0.43 | 0.30/0.70 | 1.304e-01/2.437e-03 |
| FEV<sub>1</sub> | height | 138 | 0 | 1 | 0.54/0.46 | 0.29/0.71 | 1.061e-01/9.810e-03 |
| FVC | sex | 198 | 0 | 1 | 0.51/0.49 | 0.30/0.70 | 1.037e-01/8.514e-03 |
| FVC | height | 208 | 0 | 1 | 0.52/0.48 | 0.33/0.67 | 8.071e-01/1.449e-02 |
| FEV<sub>1</sub>/FVC | sex | 266 | 0 | 0 | 0.47/0.53 | 0.40/0.60 | 1.404e-01/9.763e-03 |
| FEV<sub>1</sub>/FVC | height | 189 | 0 | 1 | 0.47/0.53 | 0.39/0.61 | 1.215e-01/1.297e-02 |
| FEV<sub>1</sub>/FVC | P-Y-S | 232 | 0 | 3 | 0.53/0.47 | 0.44/0.56 | 2.352e-02/1.283e-03 |
| FEV<sub>1</sub>/FVC | E-S | 161 | 0 | 1 | 0.55/0.45 | 0.40/0.60 | 3.771e-02/4.095e-02 |
| height | sex | 702 | 0 | 0 | 0.48/0.52 | 0.42/0.58 | 5.042e-02/3.558e-02 |

## Discussion

In this communication, we propose RITSS, a robust and flexible interaction testing framework for quantitative traits. Our framework aims to identify aggregated interaction signals and tests them using sample splitting and robust test statistics. Since interactions at the single genetic variant level are hard to detect due to small effect sizes, we hypothesize that strategies that aggregate signals across a limited number of factors/loci have higher statistical detection power while the limited number of factors/loci in the score still permit an interpretation of the aggregated signals.

In extensive simulations, we demonstrated that RITSS controls the type 1 error rates well across different scenarios, including population stratification, gene-environment correlation, mis specified environmental main effects, non-normal error distributions, and heteroscedasticity. We also applied our approach to the UK Biobank and observed potential interactions between subcomponents of risk scores and sex in lung function and height. Since interactions for complex traits at the single genetic variant level were rarely detected in recent large-scale analyses, we suggest that our approach will be an important tool for the identification of genetic interactions and the underlying mechanisms.

For example, in the context of gene-by-sex interactions, Fawcett et al. performed a genome-wide interaction analysis with sex for FEV_1_ in the UK Biobank, using more than 300,000 samples (Fawcett *et al*., 2021). Although they found five genome-wide significant interactions, only one interaction was replicated in an independent study. In a different study, Bernabeu et al. utilized the UK Biobank to analyze genotype by sex interaction for 530 complex traits. This analysis revealed several genome-wide significant (*p* < 10^−8^) findings, but heritability and genetic correlation analyses suggest that substantial proportions of the sex-differential genetic architecture are yet to be discovered (Bernabeu *et al*., 2021). Our results are in line with these findings since our interaction signals are not driven by strong effects on a single variant level but aggregations across a limited number of variants.

Our approach has the following limitations. A significant interaction p-value of the aggregated score does not imply that all included pairs of genetic variants/environmental factors necessarily truly contribute to this interaction. Depending on the specific implementation of the screening step, our approach can be computationally demanding and the size of the input set of genetic variants is restricted in size. Finally, a significant interaction p-value does not imply that causal mechanisms are detected. RITSS aims to identify deviations from the null hypothesis model 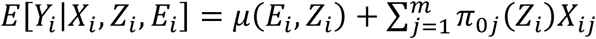 in the form of environment-dependent genetic effects *π*(*E*_*i*_). However, the interpretation of significant findings depends on the assumption that there is no unmeasured variable omitted that is involved in this mechanism (Dudbridge and Fletcher, 2014; VanderWeele, 2009).

Future directions include more detailed follow-up analyses, the extension to dichotomous traits, the incorporation of other screening techniques, such as for example LASSO (Tibshirani, 1996) or RaSE (Tian and Feng, 2021), and the improvement of the computational efficiency. We note that RITSS can also be used to analyze gene-gene-interactions and interaction models can be based on more general, non-linear approaches. In addition, RITSS can be applied to other omics data layers, such as for example, DNA methylation, transcriptomics, or metabolomics. An R package implementing RITSS and providing the framework for extensions, as well as the simulation study code, is available at https://github.com/julianhecker/RITSS.

## Appendix A

### Simulation details

The covariates *Z*_*ij*_ are simulated based on a uniform distribution *Z*_*ij*_∼*Unif*(*a, b*), *j* = 1, …, *p*. They are used to simulate population stratification (PS) and can be considered as the principal components (although not orthogonal). The parameters *a* and *b* are set to *a* = 0.0 and *b* = 0.1. The genotypes *X*_*ij*_ are generated based on a binomial distribution with *X*_*ij*_∼*Binom*(2, *p* = 0.3 + *β*_*PS*_ * *Z*_*iυ*_), *j* = 1, …, *m*, where *β*_*PS*_ = 1 in the presence of population stratification (PS:yes) and *β*_*PS*_ = 0 if not (PS:no). The index *υ* is chosen based on a varying sequence that depends on *m* and *p*, to ensure approximate coverage of all components. The environmental factors *E*_*ij*_ follow 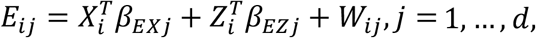 with *W*_*ij*_∼*N*(0,1). The vectors *β*_*EXj*_ and *β*_*EZj*_ control gene-environment correlations (GEC) and population differences in the environment. The residual error *ε*_*ij*_ is simulated by *ε*_*ij*_ = *b*_*ij*_ * (1 + *β*_*εE*_ * *E*_*i*1_), where *b*_*ij*_∼*N*(0,1) in the case of normal errors. In the scenario of non-normal errors (NNE), *b*_*ij*_ is sampled from the mean-centered and standardized lung function ratio in the UK Biobank (Bycroft *et al*., 2018). The parameter *β*_*εE*_ controls the presence of heteroscedastic errors (HE). The phenotype *Y*_*i*_ is then constructed by:

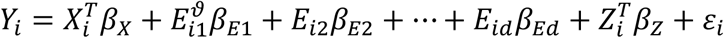

The exponent *ϑ* is introduced to simulate mis specified environmental main effects (MEM). The effects *β*_*X*_, *β*_*E*_, *β*_*Z*_, *β*_*EZj*_, *β*_*EXj*_, and *β*_*εE*_ are generated as follows:

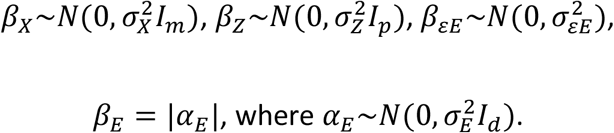

and,

*β*_*EXj*_ = *B*_*j*_|*α*_*EXj*_| where 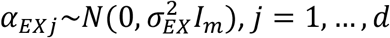 and *B*_*j*_∼*Bernoulli*(0.2) as well as

*β*_*EZj*_ = *B*′_*j*_*α*_*EZj*_ where 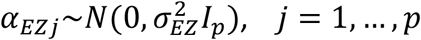 and *B*′_*j*_∼*Bernoulli*(0.5). The parameters *β*_*X*_, *β*_*E*_, *β*_*Z*_, *β*_*EZj*_, *β*_*EXj*_, and *β*_*εE*_ are freshly drawn in each replication of the simulation study. We set 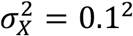 and 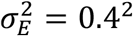. In the following Table S1, we report scenario-dependent parameter values/implementations.

**Table S1:** Scenario-dependent parameter values for the simulation studies. PS: population stratification, GEC: gene-environment correlation, MEM: mis specified environmental main effect, NNE : non-normal errors, HE : heteroscedasticity.

|  | yes | No |
| --- | --- | --- |
| PS | $\beta_{PS} = 1, \sigma_{EZ}^2 = 0.1^2, \sigma_Z^2 = 0.1^2$ | $\beta_{PS} = 0, \sigma_{EZ}^2 = 0.0, \sigma_Z^2 = 0.0$ |
| GEC | $\sigma_{EX}^2 = 0.1^2$ | $\sigma_{EX}^2 = 0.0$ |
| MEM | $\vartheta = 2$ | $\vartheta = 1$ |
| NNE | $b_{ij} \sim FEV_1/FVC \text{ UKB}$ | $b_{ij} \sim N(0,1)$ |
| HE | $\sigma_{\varepsilon E}^2 = 0.5^2$ | $\sigma_{\varepsilon E}^2 = 0.0$ |
| SELECT | $\frac{m}{2}$ variants with smallest association<br>p-value selected for analysis (based on all $n = 30,000$ samples) | keep all variants |

## Appendix B

### ACE algorithm

The input for the ACE algorithm (Breiman and Friedman, 1985; Vansteelandt *et al*., 2008) consists of the score *U*_*i*_, genotype data *X*_*i*_, environmental factors *E*_*i*_, and the covariates *Z*_*i*_. The goal is to derive a projection of *U*_*i*_ that is orthogonal to functions *μ*(*E*_*i*_, *Z*_*i*_) (i.e., 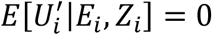) and orthogonal to the genetic main effects 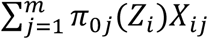.

Starting from an input score of the form *U*_*i*_ = ∑_*j*_ ∑_*l*_ *π*_*jl*_*X*_*ij*_*E*_*il*_, we

1. estimate *E*[*U*_*i*_|*X*_*i*_, *Z*_*i*_] = ∑_*l*_ *π*_*jl*_*E*[*E*_*il*_|*X*_*i*_, *Z*_*i*_] ∑_*j*_ *X*_*ij*_ using a flexible LASSO-based linear model in combination with cross-validation and replace *U*_*i*_ by the corresponding residuals.
2. Use a machine learning approach to estimate *E*[*U*_*i*_|*E*_*i*_, *Z*_*i*_] and replace *U*_*i*_ by the corresponding residuals.
3. Use a machine learning approach to estimate *E*[*U*_*i*_|*X*_*i*_, *Z*_*i*_] and replace *U*_*i*_ by the corresponding residuals.
4. Repeat steps 2.) and 3.) until a sufficient criterion is reached.

For step 2.), we applied a regression forest approach, implemented in the *grf* R package (Athey *et al*., 2019). For step 3.), we used a LASSO-based linear model with *X*_*i*_ and *Z*_*i*_ as covariates, in combination with cross validation. In practice, 2-3 repetitions of 2.) + 3.) are often sufficient.

## Appendix C

### UK Biobank data

#### Study population

Our data analysis utilized participants from the UK Biobank (Bycroft *et al*., 2018). All participants in the analysis provided written informed consent and study protocols were approved by local institutional review boards/research ethics committees. We selected participants of European ancestry only, whereas ancestry was derived based on a combination of self-reported ethnicity and k-means clustering of principal components of genetic ancestry, as previously described (Shrine *et al*., 2019). Additional quality control included the exclusion of related pairs (keeping one sample), samples with low quality lung function (Shrine *et al*., 2019; Sakornsakolpat *et al*., 2019) as well as missing phenotype or covariate data. Overall, we kept 254,053 participants for the analysis.

#### Phenotype and covariate data

We incorporated lung function data as measured by forced expiratory volume in 1 second (FEV_1_), forced vital capacity (FVC), and the ratio FEV_1_/FVC. We also extracted age, sex, height, smoking exposure variables, genotyping array, and the first ten principal components of genetic ancestry. Smoking exposure was based on self-reports and included the variables pack-years of smoking (P-Y-S) and ever-versus never-smoking status (E-S). ‘Ever-smokers’ included individuals reporting current smoking, smoking most days, or smoking occasionally. ‘Never smokers’ included those who smoked less than 100 cigarettes in their lifetime.

#### Genetic data

Genotyping and imputation for the UK Biobank were performed as described in the corresponding publications (Shrine *et al*., 2019; Bycroft *et al*., 2018) and this genetic data was available for our analyses. For each of the four phenotypes FEV_1_, FVC, FEV_1_/FVC, and height, we downloaded all reported genetic associations from the GWAS catalog (MacArthur *et al*., 2017) (February-August, 2021). We extracted the corresponding genetic variants with a minor allele frequency above 1% (estimated in the analysis dataset) and performed LD pruning *(indep-pairwise* command with parameters 500, 50, and 0.2) using PLINK2 (Version v2.00a2.3LM) (Chang *et al*., 2015) to avoid reported genetic associations that are in high LD. We also excluded multi-allelic variants. Our analyses are based on expected minor allele count information, as computed by PLINK2. The final number of variants for the analysis of the respective phenotypes is described in Table 1.

