## supplementary_figures for "A robust and adaptive framework for interaction testing in quantitative traits between multiple genetic loci and exposure variables"

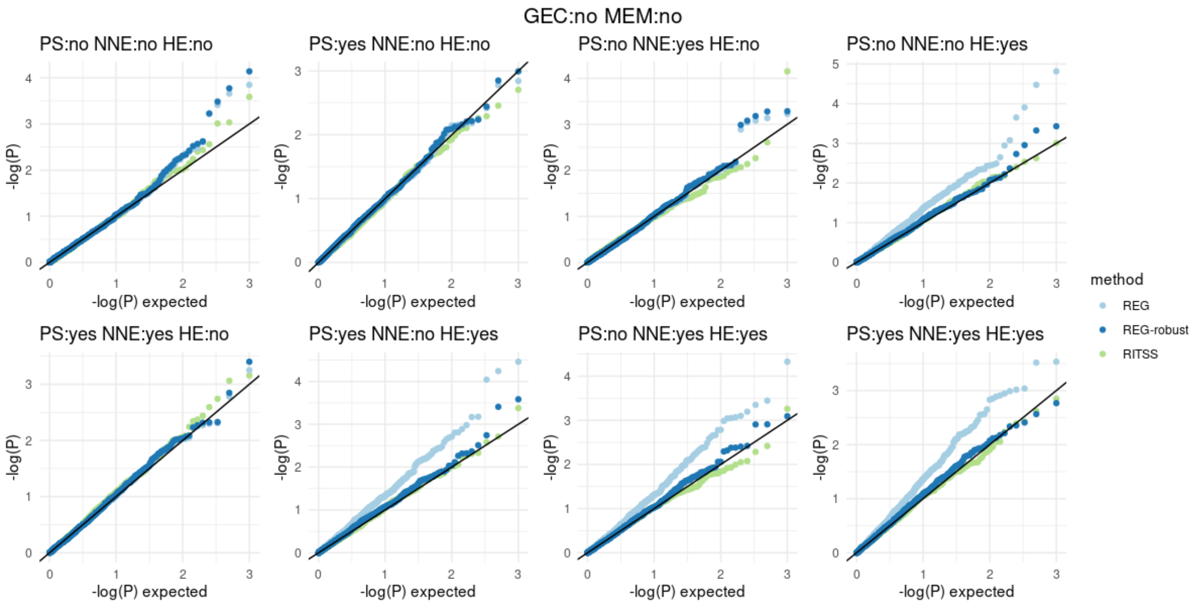


Supplementary Figure 1. Quantile-quantile-plots for RITSS, REG, and REG-robust in the 8 scenarios with SELECT:yes, GEC:no, and MEM:no. All results based on 1,000 replicates. GEC: gene-environment correlation, MEM: mis specified environmental main effect, PS: population stratification, NNE: non-normal errors, HE: heteroscedasticity.

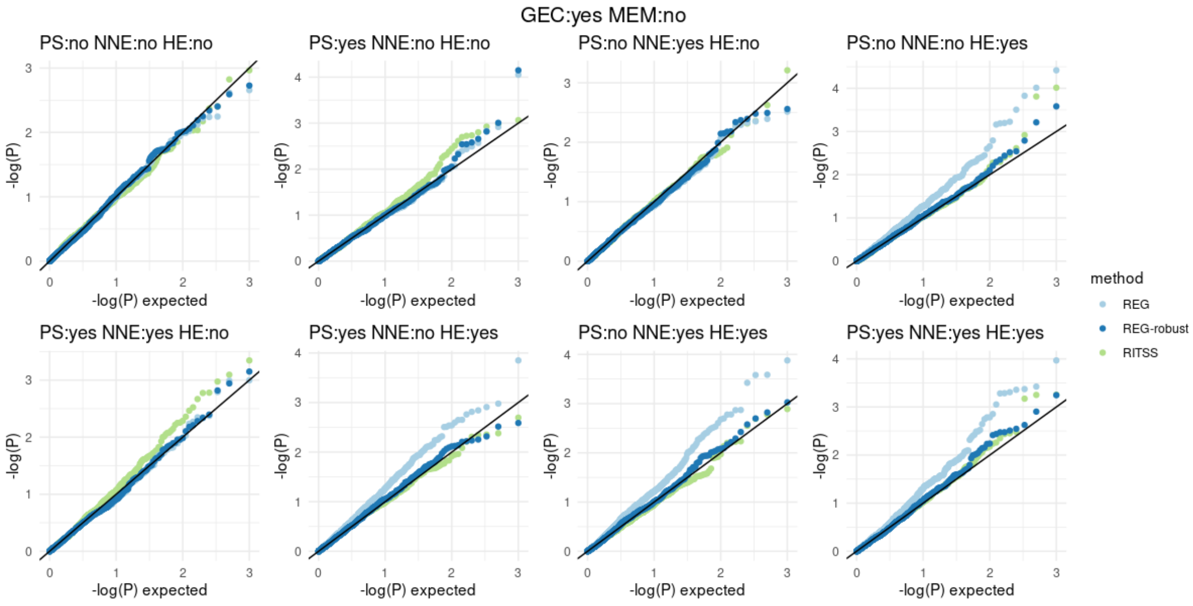


Supplementary Figure 2. Quantile-quantile-plots for RITSS, REG, and REG-robust in the 8 scenarios with SELECT:yes, GEC:yes, and MEM:no. All results based on 1,000 replicates. GEC: gene-environment correlation, MEM: mis specified environmental main effect, PS: population stratification, NNE: non-normal errors, HE: heteroscedasticity.

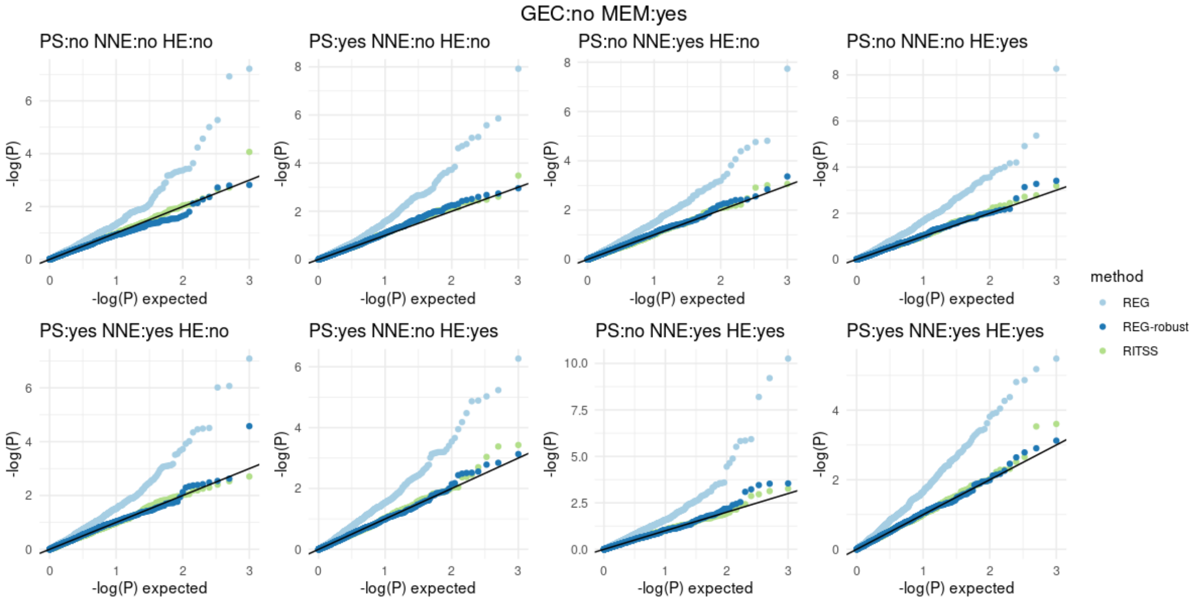


Supplementary Figure 3. Quantile-quantile-plots for RITSS, REG, and REG-robust in the 8 scenarios with SELECT:yes, GEC:no, and MEM:yes. All results based on 1,000 replicates. GEC: gene-environment correlation, MEM: mis specified environmental main effect, PS: population stratification, NNE: non-normal errors, HE: heteroscedasticity.

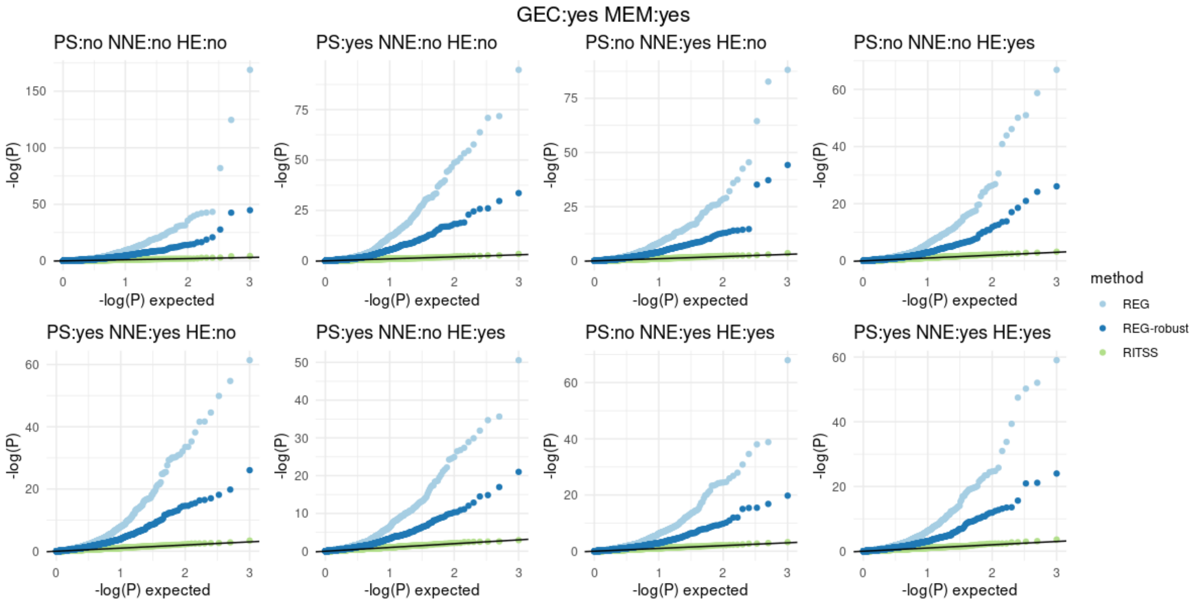


Supplementary Figure 4. Quantile-quantile-plots for RITSS, REG, and REG-robust in the 8 scenarios with SELECT:yes, GEC:yes, and MEM:yes. All results based on 1,000 replicates. GEC: gene-environment correlation, MEM: mis specified environmental main effect, PS: population stratification, NNE: non-normal errors, HE: heteroscedasticity.

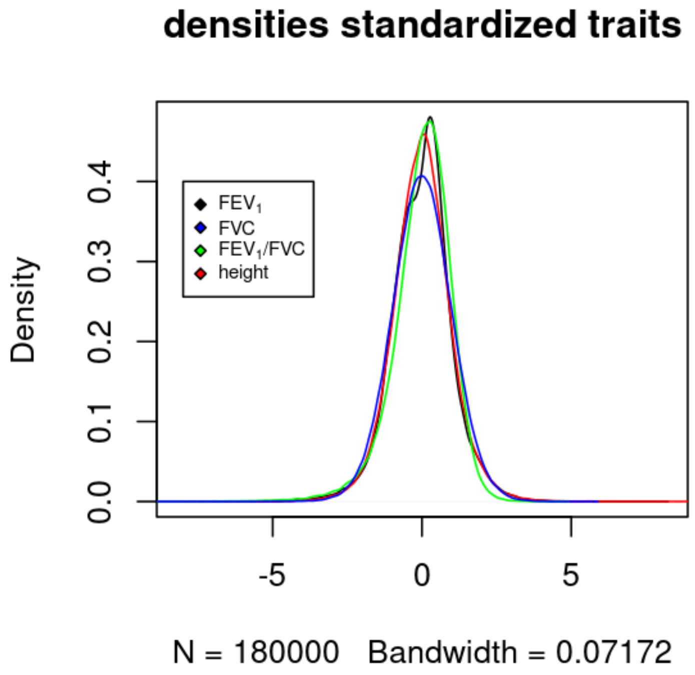


Supplementary Figure 5. Density plots for standardized residuals for all four traits in the analysis. $FEV_1$: forced expiratory volume in 1 second, FVC: forced vital capacity (FVC).
